# QuadStack: Specialized convolutional blocks enable *in vivo* BG4-binding motif prediction and highlight discrepancies with *in vitro* G-quadruplexes

**DOI:** 10.64898/2026.05.06.723235

**Authors:** Polen Nehir Ulaş, Osman Doluca

## Abstract

G-quadruplex (G4) prediction has been largely guided by *in vitro* biophysical rules, yet these models show limited agreement with *in vivo* measurements. Here, we present QuadStack, a deep learning model trained on a multi-study BG4-ChIP-seq compendium. QuadStack introduces two biologically grounded convolutional modules—G4Stack Convolution, which captures G/C stacking patterns, and Reverse Complement Convolution, which enforces strand-invariant representations consistent with ChIP-seq signals. QuadStack achieves strong predictive performance (AUC up to 0.94) and substantially outperforms widely used *in vitro*-based predictors on genomic test data. Beyond performance, our analyses reveal that BG4-associated sequence grammar is not solely governed by canonical isolated G-rich tracts, but also by patterns where G and C nucleotides are mixed. This suggests that cytosines are not simply disruptive *in vivo*, and raises the possibility that cytosines may play a context-dependent role or that guanines on the opposite strand contribute to the structure, which could explain the difference between *in vivo* and *in vitro* observations. Together these findings demonstrate a fundamental discrepancy between *in vitro* folding propensity and *in vivo* G4 biology, and establish QuadStack as both a predictive model and a framework for interpreting G4 formation in its native genomic context.

## Introduction

G-quadruplexes (G4s) are noncanonical secondary nucleic acid structures that can be present in guanine-rich sequences. Four Guanine (G) nucleotides converge on the same plane, stabilized by Hoogsteen hydrogen bonds and a monovalent cation in the center, is called a G-quartet. Coaxial stacking of G-quartets creates the G-quadruplex structure. These structures can be found in functional parts of genome like telomeres and promoter region. They play key roles in some biological processes like telomere damage signaling, DNA replication, transcription and translation regulation [14, 17]. On the other hand, G-quadruplexes can regulate oncogenes in a way that stabilization of G4 formation can inhibit cancer cell development [3]. Therefore, prediction of G-quadruplex structures has an important role in medical biology and cancer therapy as well as molecular biology.

G-quadruplexes have gained increasing attention due to their potential regulatory roles in genomic stability, transcriptional control, and therapeutic targeting, especially in cancer research. Experimental evidence has revealed that G4s are highly context-dependent, varying across cell types, physiological conditions, and chromatin states [10]. Consequently, computational prediction of G4 formation remains a challenging problem. Traditional motif-based and thermodynamics-based prediction tools are designed primarily to detect potential G4-forming sequences (PQS) and often rely on *in vitro* folding rules [12]. However, a growing body of evidence suggests that *in vivo* G4 formation is influenced by additional factors such as chromatin accessibility, transcriptional activity, local secondary structure competition, and protein binding [10, 15].

Several experimental methods have been developed to profile G4 formation genome-wide. These include G4-seq, rG4-seq, and ChIP-based approaches such as BG4-ChIP-seq, which uses the BG4 antibody to capture G4 formations *in vivo*. Among these, BG4-ChIP-seq provides direct evidence of G4 binding in chromatin, reflecting physiologically relevant formation states [7]. Yet, interpreting BG4 signals and predicting *in vivo* G4 formation from sequence alone remains nontrivial.

Deep learning models have recently been applied to G4 prediction due to their ability to learn complex sequence patterns. Convolutional neural networks (CNNs) and recurrent architectures have been trained on G4-seq or *in vitro* datasets to predict PQS and G4 folding potential. While these models can achieve high accuracy within a given dataset, they often fail to generalize across species or experimental contexts, suggesting that they may not capture the core biological constraints driving BG4-binding and *in vivo* G4 formation.

In this work, we hypothesize that capturing *in vivo* BG4-associated signals requires models that explicitly encode key biological rules of G4s and BG4-ChIP-seq behavior. In particular, two such rules are (i) the dominance of guanine stacking and G/C balance in stable G4 contexts and (ii) the strand-invariant nature of BG4-ChIP-seq binding, where the antibody binds folded structures irrespective of genomic strand annotation.

To address these gaps, we focus explicitly on *in vivo* BG4-associated G4 biology by training on an extensive, multi-study BG4-ChIP-seq collection, enabling a first large-scale *in vivo* sequence model of BG4-binding. In doing so, we (a) identify two novel convolutional blocks that may generalize to future DNA sequence models. Finally, by contrasting BG4-positive loci with *in vitro*-oriented predictors and control sets, we (b) highlight major *in vivo*–*in vitro* differences, (c) provide evidence that complementary-strand motifs can contribute to *in vivo* G4 behavior, and (d) cytosines are not necessarily inhibitory in the genomic DNA context.

## Methods

### Data

Datasets were generated based on high-throughput G4-ChIP-seq studies using BG4 antibody available at NCBI GEO Gene Expression Omnibus, GEO Datasets. Only the experiments conducted between 2016 and 2024 were considered for analysis. Eight distinct studies employing BG4 were utilized to generate datasets, each identified by the GSE (GEO Series) accession codes GSE76688, GSE99205, GSE107690, GSE145090, GSE161531, GSE162299, GSE225772, and GSE241008. For each GSM (GEO Sample), raw data of technical replicates and input (control) data were accessed by NCBI’s Sequence Read Archive (SRA) Run code. (see Supplementary Information)

### Galaxy workflow

Raw data that have been utilized in this study were processed by using Galaxy Bioinformatics web server available at https://usegalaxy.org/ which provides computational tools for data upload, processing and alignment. From galaxy home, Tools section was opened and Get Data tool was used [16]. The Download and Extract Reads in FASTQ function of the tool was utilized to retrieve and store the raw sequencing reads in fastq format, ensuring compatibility for downstream analysis. The input type was selected as “SRR accession” and SRR accession code of each technical replicate was entered to the relevant text box. “Gzip compressed fastq” was selected as the output format before run. Build List function under Collection Operations was used to group technical replicates of the same biological sample in order to pool them in the peak-calling step. The names of the grouped lists were edited with their biological replicate and technical replicate number as well as whether they served as input.

Individual Galaxy tools linked in a way that the output of one tool serves as the input for the subsequent, enabling the automation of multi-step analysis and ensures that all data undergo the same processes. The phases of the G4-ChIP-seq data analysis workflow are explained in the supplementary Figure S1 and the following:

1. The treatment files grouped by the build list function were submitted to the first step of the workflow as input. In the same manner, the control file was submitted to the workflow as another input.
2. The treatment files went through the **FASTQC** analysis tool (Galaxy Version 0.74+galaxy1) in order to analyze the quality of the raw data read and adapter content. All parameters were set to default.
3. Depending on the results of FASTQC analysis, both treatment and control files in fastq format underwent a trimming process individually by the **Trimmomatic** tool (Galaxy Version 0.39+galaxy2), which enables flexible read trimming for Illumina NGS data. For tool parameters, the data type was set to single-end for single-end data and paired-end for paired-end data. `Perform initial ILLUMINACLIP step` was set to *yes*. Standard adapter sequences were chosen for trimming and appropriate adapter sequences were used; other adapter parameters were set to default. Overall, trimmomatic operations were applied using the following parameters, `LEADING`: 3, `TRAILING`: 3, `SLIDINGWINDOW`: 4:20.
4. Both treatment and control data were evaluated by FASTQC analysis after the trimming step to confirm the removal of adapter sequences.
5. The trimmed treatment and control data were aligned separately on a reference genome using **Bowtie2** (Galaxy Version 2.5.3+galaxy1). Single-end library was selected as the library parameter. A built-in genome index was used and *Human (Homo sapiens) (hg38): hg38 Canonical* was selected as the reference genome. As the `preset`, `very-sensitive end-to-end (--very-sensitive`) parameter was applied. Other parameters were left as default. The outputs of the **Bowtie2** were received as BAM files.
6. To sort treatment and control BAM files independently, the **Samtools sort** tool (Galaxy Version 2.0.5) was used with all parameters set to default.
7. Duplicate sequences were marked using the **MarkDuplicates** tool (Galaxy Version 3.1.1.0) after sorting by setting the `assume the input file is already sorted` parameter to *yes* and *SUM OF BASE QUALITIES* was selected as the scoring strategy for the `choosing the non-duplicate among candidates` parameter.
8. Treatment and control BAM files followed the peak calling process by the **MACS2 callpeak** tool (Galaxy Version 2.2.9.1+galaxy0). `Pooling files` and `control file` parameters were set to *yes*. The `effective genome size` was set as *H. sapiens* (2.7 × 10^9^). The outputs of the tool were received as `output narrowpeaks` and `output summits` in bed format.

### Preprocessing

The output files were retrieved from the Galaxy server in Narrowpeak format, where the columns include chromosome number/name, start position of the peak, end position of the peak, peak name, score, strand, signal value, p-value, q-value and peak position (summit) relative to start position of the peak, respectively. The peaks had variable-length sequences and since neural networks with fully connected layers can receive only fixed-length inputs, peak intervals were divided into fixed-length windows as followed. Starting from 50 bases upstream of the peak position, 100 bases were extracted and repeated as the window shifts by 50 bases downstream until the peak end is reached. In the same manner, backwards extraction started at 100 bases upstream of the peak position and shifted by 50 bases upstream each time until interval start was reached. The extracted sequences were added as a new column to the DataFrame, and saved (.a files).

Finally, all of the sequences were arranged such that each row contains the chromosome number and the 100 nt-long sequence respectively and colon-separated. This format for all positive and negative control inputs of the study were maintained and stored as .c files. Prior to loading the model, each positive and negative sequences were doubled by reverse complementing.

### Negative controls (NC)

#### Randomly shifted negatives

To generate negatives matched to local genomic context, we constructed a randomly shifted negative-control (NC) set. For each positive peak, we extracted a window of identical length after shifting the peak center by a random offset sampled uniformly from [−10*kb*, +10*kb*] bp. Candidate shifted windows were discarded if they overlapped any positive peak in the union of all BG4 datasets used in this study. This procedure yields negatives that are proximal to the positives and therefore approximately matched for local genomic (and putatively chromatin) context, while avoiding direct reuse of labeled positive regions.

#### Shuffled negatives

Shuffled negative controls (NCs) were used to evaluate whether models learn discriminatory features beyond simple base composition. For each positive sequence, we generated a negative example by shuffling nucleotides while preserving sequence length (and, where applicable, nucleotide or dinucleotide composition). This procedure retains global compositional properties (e.g., G/GC enrichment) but disrupts positional organization of G-runs and loop-like spacing patterns required for G4 formation. Performance against shuffled NCs therefore provides a stringent test of whether the model captures higher-order sequence organization rather than relying on trivial compositional cues.

#### Non-BG4-binding quadruplexes

Finally, computationally obtained G-quadruplex predictions from the human genome was compared to the positive dataset. G4Catchall was run using the standard *G3+E3+I1B* parameter configuration[5]. This computational list was then compared against the BG4-ChIP-seq positive coordinates for overlap detection. Predicted non-overlapping sequences were extracted as the final negative control dataset. All negative control data were also processed and converted to .c file where necessary, maintaining the previously described format.

#### Data separation

From both positive data and negative control, except shuffled NC, any sequence obtained through chromosome 19 and chromosome 20 sequences were reserved for testing and validation, respectively, so no sequence overlap would exist between training, validation, and test sets. The number of sequences used for each set is detailed in Table S1. Equal number of positive and negative chromosome 19 data was used to create the test set, ensuring that any untrained model would deliver 0.5 precision. During training, the validation set (chromosome 20) was monitored to trigger early stopping and learning rate adjustments. To evaluate the results, Area under the ROC (Receiver Operating Characteristic) curve and AP (Average Precision) score were utilized. The ROC and PR curve were plotted for each session.

#### Model architecture

Two types of inputs were supplied to the model, the 100-nt long sequences in one-hot encoded format, with one base per column. The model was built to generate a binary classification indicating whether the given sequence is classified as positive or negative for BG4-binding. To test the different model architectures, the information was either directly passed to a core convolution neural network (CNN) block or two novel convolution architectures, G4Stack Convolution (G4SC) and Reverse Complement Convolution Blocks (RCC), before passing to the core block. The binary cross-entropy was chosen as the loss function and AUC-ROC (Area under the receiver operating characteristic curve) and AUC-PR (Area under the precision-recall curve) as the metric. The optimizer was selected as Adam.

#### G4Stack Convolution (G4SC) Block

Since the G4 formation is considered necessary for BG4 to bind, the weight of Guanines and Cytosines are expected to be significantly higher than other nucleotides. For that reason, the model can benefit from evaluating Guanine and Cytosine channels preliminarily. To do so, a new G4Stack Convolution Block was designed, specifically computing on these two channels. Each Guanine and Cytosine channel was extracted and a 1D convolutional layer was applied. To capture varying lengths of G-tracts, three different kernel sizes (3, 5, and 7) were applied. The outputs were concatenated to the original input, creating new tensors with (100,11) dimensions (Figure 2a).

**Fig. 1.**
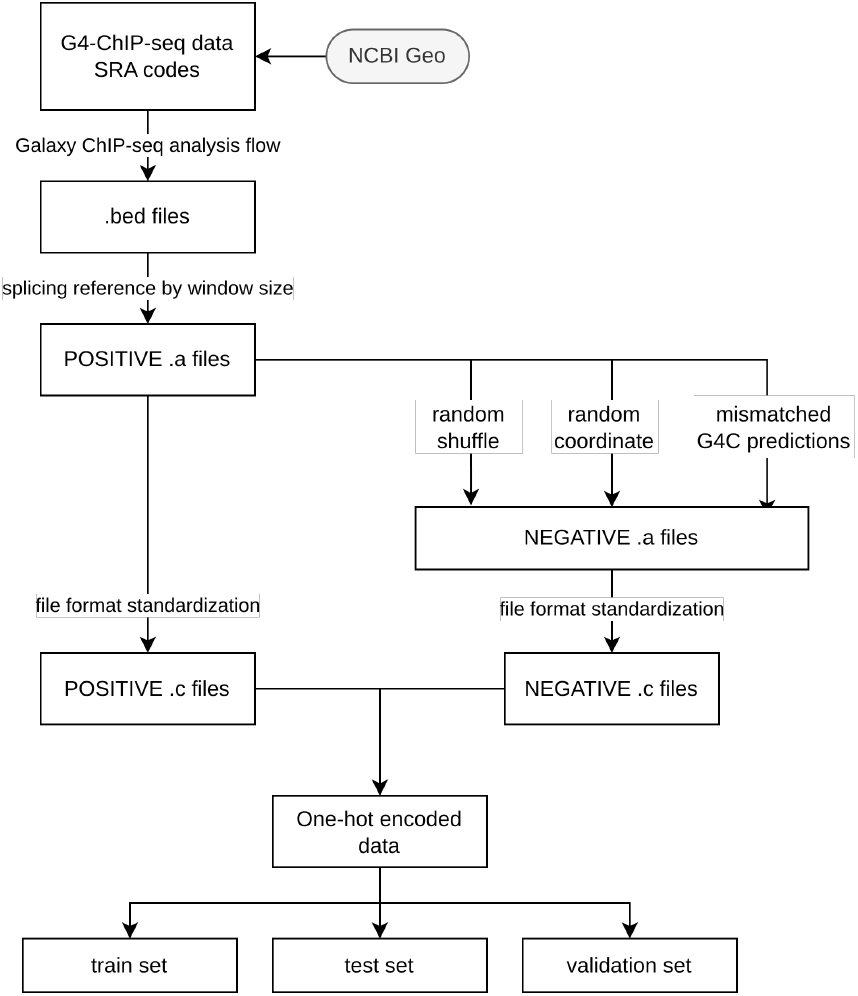
Overview of data flow in this study.

**Fig. 2.**
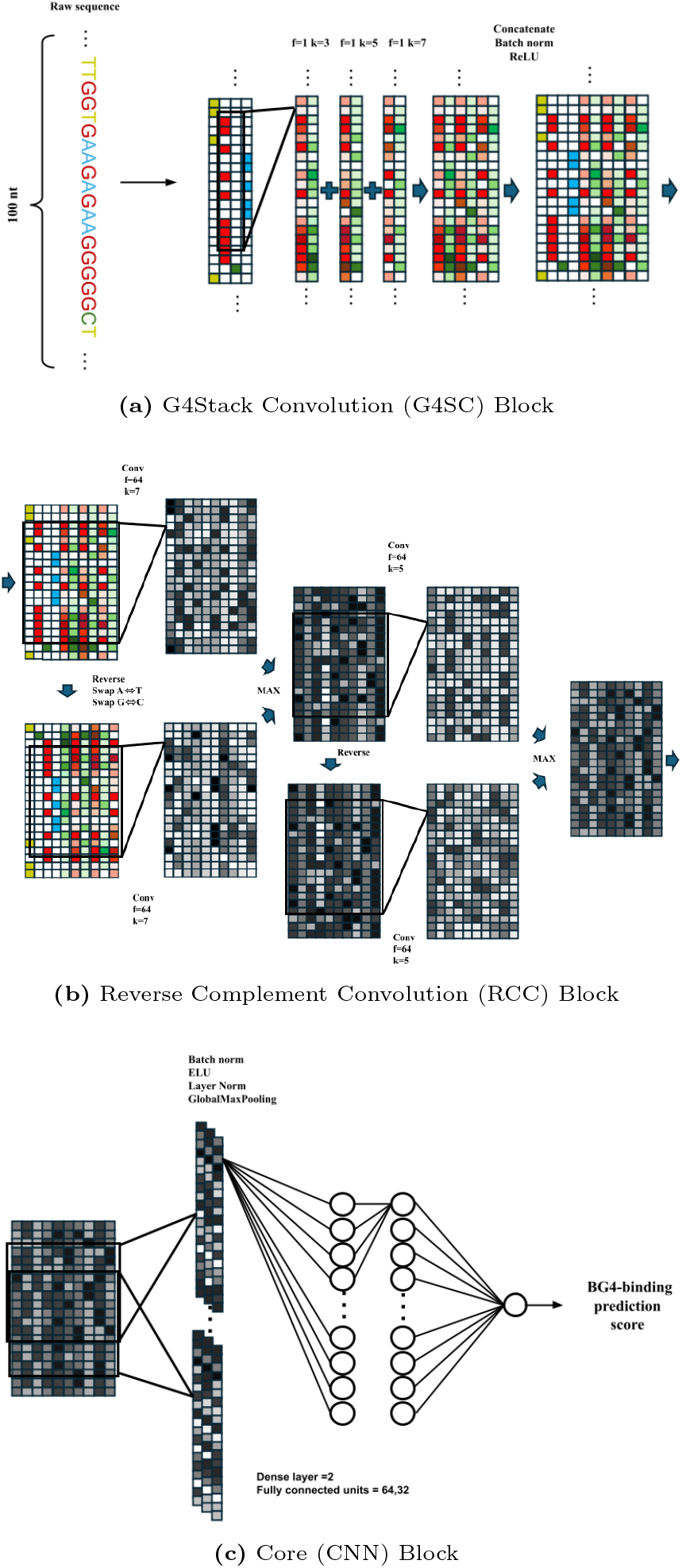
The overall architecture of the model, detailing its constituent blocks.

#### Reverse Complement Convolution (RCC) Block

This model is aimed at predicting BG4-binding based on BG4-ChIP-seq, which yields both strands, even if the G-quadruplex is formed only on one. For this reason, Reverse Complement Convolution Block was added to train the model with reverse complementary sequence and probability of the original input data. The output taken from G4SC Block was processed as follows. Initially the input was reversed in the sequence axis. This was followed by swapping Adenine channel with Thymine’s, Guanine channel with Cytosine’s and any G4SC obtained from Guanine channel with consecutive G4SC channel obtained from Cytosine channel. Next, a convolutional layer (filters=64, kernels=7) was applied to both the reversed and the original input. The maximum of each position was taken to form an output. This process was repeated using the output for a second time, except without swapping the channels again (filters=64, kernels=5) (Figure 2b).

#### Core CNN Block

The core CNN block was composed of two convolutional layers which were joined end-to-end and two dense layers. All convolutional layers have ELU (Exponential Linear Unit) as activation function to provide nonlinearity and layer normalization by applying normalization statistics to outputs of layers in order to maintain internal stability. Also, batch normalization was added to prevent overfitting and maintain external stability. The design continued with GlobalMaxPooling1D to capture the most prominent feature across the entire sequence dimension, and dense layers to demonstrate effective feature extraction performance. Dense layers also used ELU as the activation function except the output layer which used sigmoid activation function in order to distinguish binary classification (Figure 2c). The overall architecture of the model is represented in (Figure 2).

#### Block Assembly

Four different model architectures were built, as the combinations of G4SC, and RCC blocks. The largest model receives the input into the G4SC Block to highlight the Guanine and Cytosine signals (Figure 2a), followed by Reverse Complement Convolution Block to introduce strand-invariance. Maximum feature of each strand is taken (Figure 2b). Resulting feature map is passed through the Core Block which includes global max-pooling layer and two fully connected layers. The output is executed by a single neuron with sigmoid activation to predict the G4-forming score of the input (Figure 2c). Three smaller models were built without G4SC, RCC or both.

## Results and Discussion

Recent evaluations of G4-ChIP-Seq data have revealed considerable heterogeneity in peak calls, with only a minority of peaks consistently shared across all biological replicates within individual studies [18]. Because intracellular G4 formation is a highly dynamic process influenced by transient cellular activities such as transcription and replication, a single-study would highly be susceptible to technical and biological noise.

By harmonizing 35 biological samples across 8 studies, we overcome this inter-replicate inconsistency. (see Supplementary Information)

### Hyperparameter optimization

#### Optimizing Core CNN Block

A validation set, generated from chromosome 20 was used to optimize hyperparameters for the CNN Block, followed by ANOVA test in order to uncover the significance of the selected hyperparameters. The calculated p-values for each parameter indicated that only the kernel sizes of each layer have a significant impact on the overall model performance (Table S2, Figure S2). Specifically, *k*_1_ = 5 consistently yielded high performance and it’s the combination with *k*_2_ = 13 was selected as the optimal architecture. Varying filter sizes did not show improvement (Table S2) indicating that the model does not require a large number of filters to sufficiently detect G4-associated motifs. The optimal number of filters was achieved using 16 and 40, respectively. For the following fully connected layers, the optimal configuration consisted of 64, 32 units, with marginal difference between tested unit sizes. Learning rate was scheduled with `ReduceLROnPlateau` to reduce the learning rate automatically when the model’s predictive ability stops improving during training (plateau). The initial learning rate was set to 1 × 10^−6^ with Adam optimizer. The model was configured with a maximum training limit of 20 epochs. However, `EarlyStopping` callback stops the training in between 8 and 15 epochs showing that the model achieved its optimal generalization capacity.

#### G4Stack Convolution (G4SC) Block

Integration of the G4Stack Convolution (G4SC) Block led to a measurable improvement in predictive performance, reflected on AUC-ROC evaluation. For the model with no novel convolutional blocks, the mean AUC-ROC of 16 trials were computed as 0.892. The AUC-ROC value of the model with only addition of G4SC Block was averaged to be 0.918 (Figure 3). G-quadruplexes are especially prominent when guanines are stacked and consecutively stabilize the G-quadruplexes [11]. In alignment with that, the model benefitted from evaluating guanine and cytosine channels preliminarily.

**Fig. 3.**
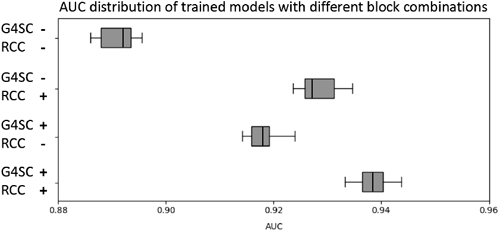
Boxplot visualization of ANOVA test results, comparing the performance distributions of various model architectures combining G4SC, RCC blocks.

**Fig. 4.**
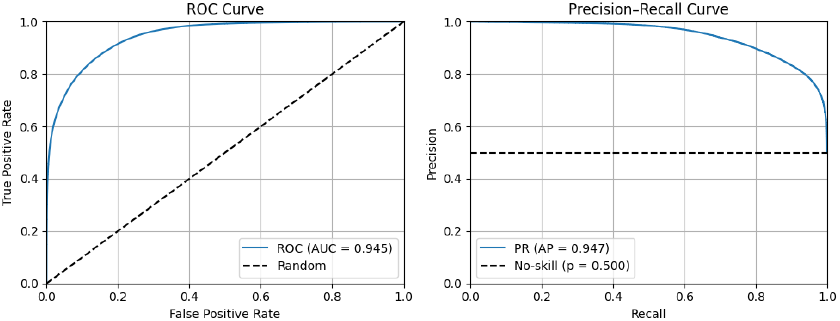
Optimal performing model achieved a peak AUC-ROC of 0.945 and AP of 0.974, significantly outperforming other architectures.

#### Reverse Complement Convolution (RCC) Block

Implementing Reverse Complement Convolution Block alone improved the model’s ability to detect G4 motifs with a mean AUC-ROC of 0.936 (Figure 3). Symmetry of the input data was provided by taking the maximum of forward and reverse feature maps, with the aim of the model to keep the strongest useful signal regardless of the sequence direction. ANOVA statistics revealed that *p*-value for contribution of RCC on overall AUC-ROC variance is 1.99 × 10^−4^, representing the block as an influential architectural component. (Table S3)

#### Synergistic Effect of Novel CNN Blocks

The mean AUC-ROC values for models with only core architecture, RCC, G4SC and G4SC:RCC were calculated from 16 different trials for each configuration (Figure 3). Amplification of G/C-derived signals by G4SC and generalizing these enriched patterns to both orientations by RCC results in the best predictive ability of the model. The average AUC-ROC of the combined model was calculated as 0.939 and the optimal model achieved 0.943 AUC-ROC and 0.947 AP.

#### Cross-Species Generalization

While at the time of the preparation of this manuscript, the majority of the available data is based on human cell lines, two studies are found on the BG4-ChIP-seq in mouse and rice genomes. To visualize the capacity of this model on both human and non-human species, we have compared the predictive accuracy across different thresholds (0.0 to 1.0) using mouse (*Mus musculus*) and rice (*Oryza sativa*), as well as human positive and negative datasets generated from chromosome 19 where applicable (Figure 5, Table 1).

**Table 1.** Comprehensive performance metrics for species-specific datasets paired with Random Shift NC controls. Optimal threshold represents the point of maximum Matthews Correlation (MCC).

| Species | AUC-ROC | PR-AUC | Max MCC | Opt. Threshold |
| --- | --- | --- | --- | --- |
| Human | 0.915 | 0.922 | 0.659 | 0.60 |
| Mouse | 0.704 | 0.944 | 0.247 | 0.10 |
| Rice | 0.578 | 0.986 | 0.048 | 0.05 |

**Fig. 5.**
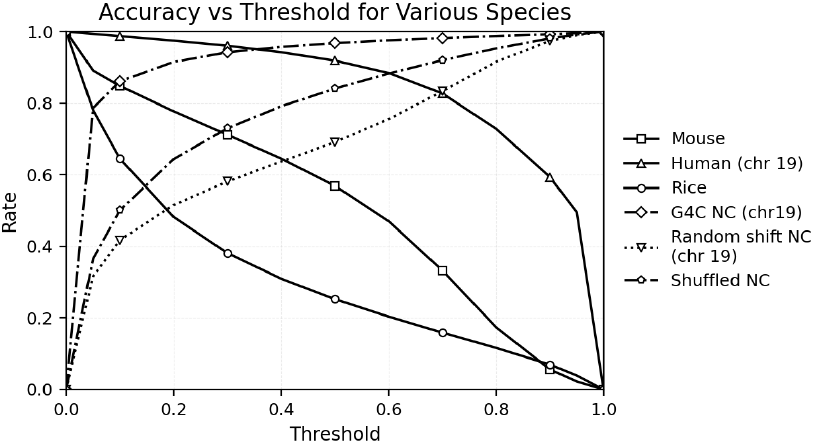
Performance of the optimal combined (G4SC:RCC) model on unseen positive test data of human (chr 19), mouse, rice, and negative datasets. For positive test data, the Y-axis represents Recall (Sensitivity), the BG4-binding sequences that model correctly predicts. In case of the negative controls represents, the Y-axis represents Specificity (True Negative Rate), the non-BG4-binding sequences that model correctly predicts.

The predictive power of human chromosome 19 (the positive test set) indicated high confidence in human-specific generalization. On the other hand, The model shows significantly less predictive power for mouse and rice data, gradually. The AUC-ROC scores demonstrate that the model can identify the conserved mammalian G4 motifs across humans and mouse confidently (0.915 and 0.704, respectively), when paired with Random shift NC (Table 1). Conversely, rice data has a sharp drop of performance at much lower thresholds, and a AUC-ROC score close to 0.578, which corresponds to marginal predictive power, indicating the model is not successful at generalizing plant genome. This failure suggests the diversity of G4-forming motifs between mammalian and plant genome may be explained by the divergence in G-richness and sequence composition of the mammalian and plant lineages [13]. It is important to note that both mouse and rice data originates from single studies, and evaluation using future studies can improve the evaluation.

In case of negative data sets, the specificity of the model against Random shift NC (chr19) and Shuffled NC negative datasets gradually increases with the threshold. Interestingly, the model showed a very high specificity towards putative G4, predicted by the algorithmic G4Catchall tool, and yet not detected in the BG4-ChIP-seq analyzes (G4C NC (chr19)). This showed that the model has learned to identify the G4 patterns that would not be bound by BG4, whether these putative G4s are biologically relevant or not.

#### In vitro vs In vivo G-quadruplex Discrepancy

As mentioned above, the model specializes in only BG4-binding motifs and learned to ignore PQS motifs that do not bind BG4 by the inclusion of putative G4s predicted by G4Catchall as part of the negatives. To benchmark the skipping of predicted G4s, we utilized another G4 prediction tool, G4Hunter [2]. G4Hunter returns a single score ranging between 4 and -4, where the sign indicates which strand has higher capacity to form G4—positive values indicating higher probability of + strand to form a G-quadruplex and vice versa. It must be noted that because, guanines on opposite strands tend to decrease the G4Hunter score, it cannot evaluate an inter-strand G4 formation. Since BG4-ChIP-seq data lacks strand discrimination, the absolute magnitude of the G4Hunter score was used to evaluate the test dataset. We also evaluated our model against a previously developed deep learning model, trained using *in vitro* G-quadruplex structures, G4Detector [1]. These tools were evaluated using their continuous scoring outputs to calculate threshold-independent metrics (AUC-ROC and AP), ensuring that performance was not limited by predefined classification cutoffs.

When evaluated on the test data originated from chromosome 19, both G4Hunter and G4Detector show significantly reduced predictive performance (AUC=0.55 for G4H, AUC=0.58 for G4Detector with K^+^ and AUC=0.57 for G4Detector with K^+^+PDS, Figure 6).

**Fig. 6.**
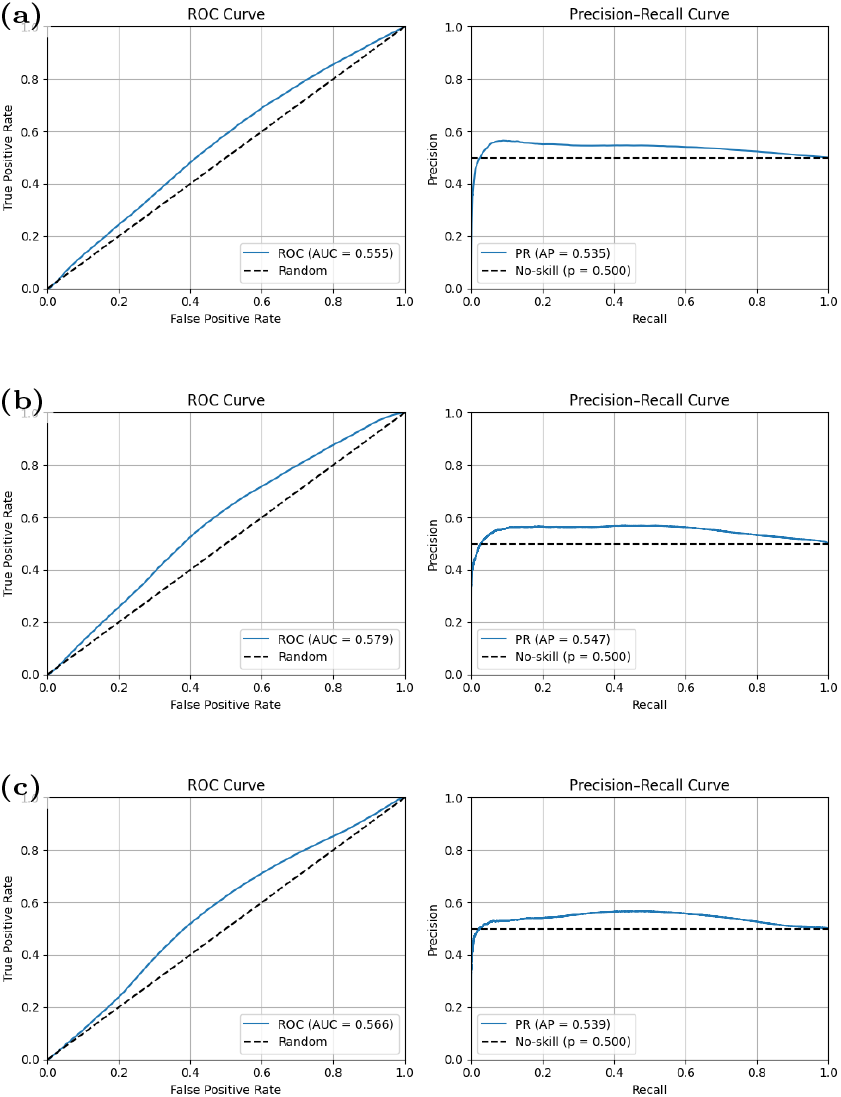
ROC and AP curves for G4Hunter (a) and G4Detector models based on K^+^ (b) and K^+^ + PDS (c) and using Human Chromsome 19 test dataset.

The performance gap between QuadStack and the existing G4-predicting tools highlights a fundamental difference in how genomic context is handled, revealing a clear divergence between *in vitro* folding propensity and *in vivo* BG4-binding. These observations suggest that *in vivo* G4 behavior is governed by additional constraints largely absent from existing predictors. The inability of *in vitro*-trained model, G4Detector, to generalize to our *in vivo* benchmark likely stems from the simplified nature of *in vitro* assays, which rely on truncated constructs lacking native flanking sequences, complementary bases or supercoiling effect resulting in biased structural outcomes. Additionally, G4 stability in a genomic setting is strongly influenced by the cellular microenvironment, such as G4-binding proteins [18, 15, 17, 7]. Many sequences predicted as G4-forming by *in vitro*-oriented tools are not enriched among BG4-positive loci, while BG4-positive regions are not fully captured by canonical PQS rules. In the following analyses, we test this interpretation by examining sequence and structural signals in a genomic DNA context.

Canonical motifs vs model

To characterize the features learned by QuadStack, we evaluated the model on a panel of artificially generated sequences derived from canonical G-quadruplex rules, systematically varying G-tract length (2-4) and loop length (1-8). Each motif was randomly placed in random 100-nt long AT-rich stretches, making an artificial benchmark dataset of 2400 sequences. When analyzed using the optimum model, we observed that among canonical PQS sequences, *G*_2_*N*_3_*G*_2_*N*_3_*G*_2_*N*_3_*G*_2_ (G_2_N_3_) was clearly more preferable. (Fig. 7)

**Fig. 7.**
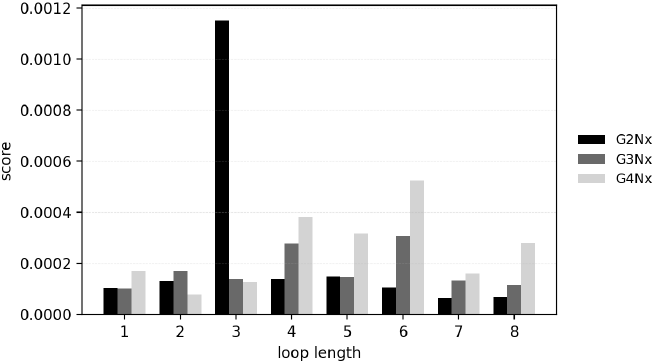
Scores of sequences based on canonical PQS rules with varying G-tract and loop lengths.

To directly evaluate nucleotide contributions, we performed systematic mutagenesis by substituting guanines inside the G-tracts of these canonical PQS sequenes with other nucleotides. Swapping guanines with A and T consistently decreased the mean score, and this trend continued until all guanines were replaced. (Fig. 8a) Surprisingly, with every substitution with cystosine increased the mean score, initially dramatically and peaking when half of the guanines were swapped. In fact swapping all guanines with cytosines were made marginal difference. If the model had simply ignored the difference between C and G, the substitutions would not have caused a change in the score, but instead it preferred a close to equal mixture of them. This alone, of course, doesn’t reveal the pattern that the model prefers, since these score are relatively quite low and far from the patterns for BG4-binding.

**Fig. 8.**
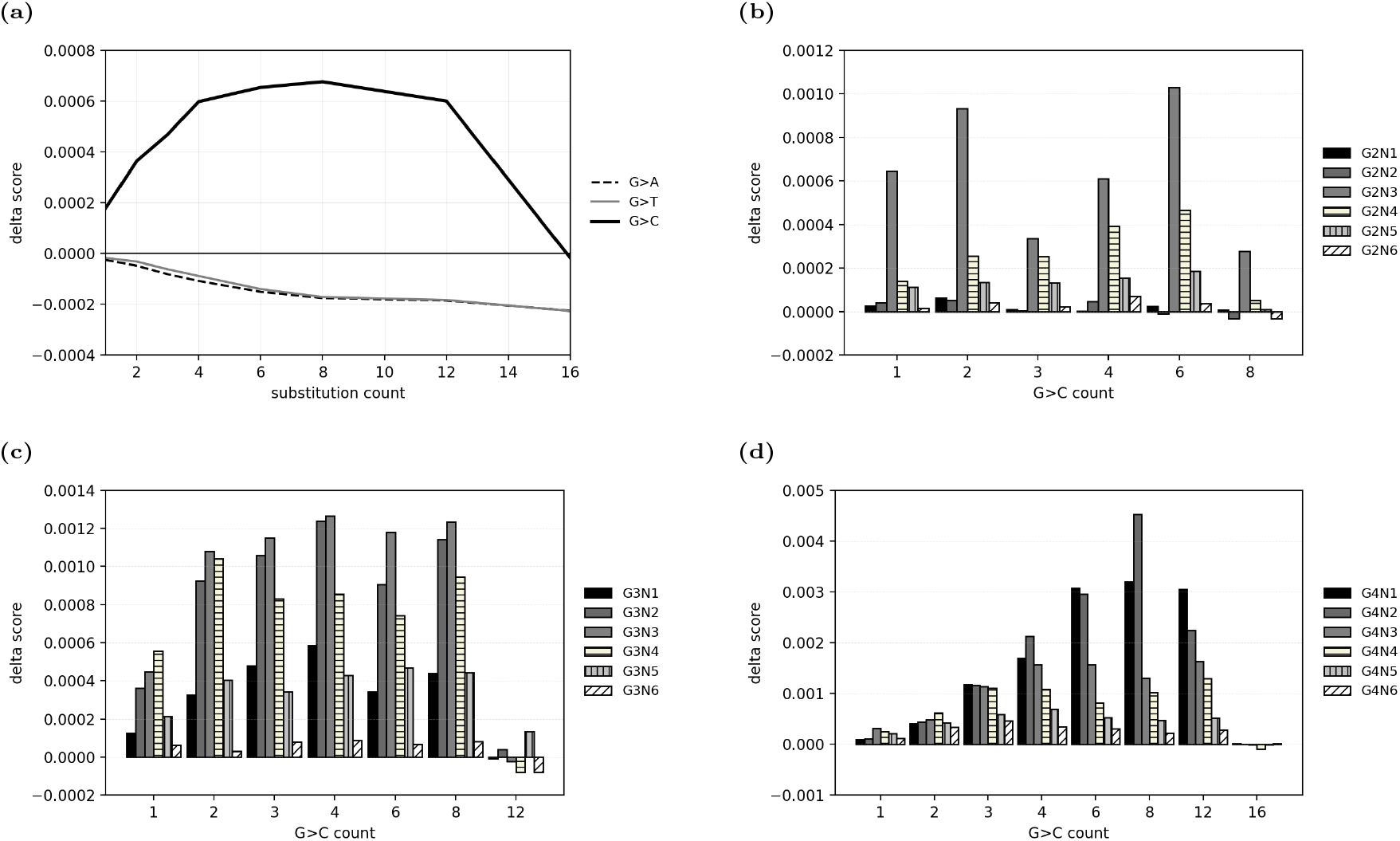
Overall effect of tract mutation count on score (a) and the impact of *G > C* substitution on score for G_2_N_x_ (b), G_3_N_x_ (c), G_4_N_x_ (d) motifs.

To assert that this was not due to asymmetric strand preference, we have also compared random sequences versus their complementary. The resulting predictions were effectively invariant (mean absolute difference *≈* 9.3 × 10^−7^), confirming strand-symmetric behavior.

To further investigate the preference G/C balance, we have observed the impact of substitutions per canonical motif (Figure 8b-d). When compared by the loop length, there was no clear preference in substitution count for G_3_N_x_ and G_2_N_x_ tracts. For both tract-lengths, 2 and 3, the score did not peak at 50% of the guanine substitutions, unlike how it was observed overall (Figure 8a). On the other hand, this was not the case for G_4_N_x_, where a clear preference for 50% (8 out of 16) of all possible substitutions persists. Observed score changes exceeded those from random perturbations with equal GC content (*p >* 0.05). Finally, another observation from these analysis was that these substitutions were more effective when the loop lengths short (1 or 2) for G_4_N_x_.

Considering together with the observation that G_2_N_3_ were the most preferred canonical PQS, and substituting guanines in G_4_N_1_ and G_4_N_2_ with cytosines had largest increase in the score, it starts to show a pattern of what the model prefers. Since cytosines do not contribute the quartet formation, substituting guanines at the end of the tracts would create longer loops fringed with cytosines and shorter G-tracts. In terms of canonical PQS rules, it is more likely that the model looks for short G-tracts with cytosines in the loop. Next, we decided to look at this preference from a different perspective. In the following section we try to identify how the highly scoring sequences from our test dataset are composed.

### The “Cytosine Myth”

It has been assumed that cytosines in a motif negatively influences the G-quadruplex formation. Yet, our results indicate that mixed C/G-rich sequence environments can still be strong BG4 targets *in vivo*. The results suggested the loops can benefit from cytosine presence and that the genomic context alters the effective grammar of BG4-associated G4s relative to purified single-strand *in vitro* folding rules. A plausible explanation is that complementary-strand sequence features can support quadruplex formation including through inter-strand guanine contribution to G-quadruplex formation, making the suggestion that presence of cytosines and guanines along the same strand causing destabilization is an oversimplification of thermodynamic preference. We investigate this hypothesis below using attribution and sequence-composition analyses on high scored stretches of sequences learned by the model rather than synthetic canonical PQS.

We analyzed 8667 sequences from the positive Human Chromosome 19 test set that scored above 0.95. The average GC content of these sequences were calculated as 0.67. The G/C balance of each sequence was evaluated by calculating *log*(*G/C*) and found to be close to zero (−0.0013), however, with a standard deviation of 0.256, well above the expected 0.109 for random sequences. This indicates that the majority of sequences has Guanine accumulation in one or the other strand, supporting that structural features are being detected by the model. Similar observation is done with the analysis of CpG density in these sequences, which reveals a strong and consistent depletion of CpG sites compared to what would be expected by random chance (0.077 vs 0.104). This means these sequences contain about 26% fewer CpG sites than a random shuffle of the same bases would produce. When these sequences were analyzed for k-mers (k=3,7,15,21), expectedly, CCC and GGG are found to be the most frequent 3-mers, followed by GGC and GCC (frequencies= 0.050, 0.052, 0.050, 0.052, respectively), well above the expected frequency of 0.016 and in alignment with our previous observations (see Supplementary Information). Infact, this observation was also made previously among high-confidence G4s (G4-II) [6]. Among 7-mers, CCCGCCC, GGGCGGG, CCCCGCC, GGCGGGG, GGGGCGG, CCGCCCC showed dramatically increased frequencies (0.0043, 0.0043, 0.0042, 0.0041, respectively).

Collectively, these observations support a revised view of BG4-associated *in vivo* G4 sequence context: enrichment of both G- and C-tracts among high-confidence positives, together with near-zero *log*(*G/C*) on average but elevated dispersion, is consistent with strand-asymmetric G-richness distributed across the two strands rather than a purely single-strand PQS rule. In this setting, cytosines can appear prominently in BG4-positive loci because they mark complementary-strand guanine tracts and/or participate in local duplex-context constraints that shape formation and recognition, explaining why destabilizing assumption of cytosines can fail as a blanket assumption outside purified single-strand *in vitro* folding. The concomitant CpG depletion further indicates that these loci are not simply explained by generic GC-richness, but reflect a more specific sequence organization relevant to *in vivo* BG4-binding.

#### BG4-binding Motifs in Chromosome 19

Identification of individual BG4-binding motifs is beyond the scope of this study. However, in order to demonstrate, De Bruijn graph assembly was applied to the identify conserved sequences using discovered k-mers. The assembled contigs were then annotated on chromosome 19, revealing a number of BG4-binding loci (Table 2). Majority of the contigs were identified to be located at the promoter regions for protein coding genes, reflecting their biological relevance. A targeted literature search did not identify prior direct experimental reports of G-quadruplex formation nearby most of the genes highlighted here; except, indications of G-quadruplex-associated regulation for AP2A1, DMPK and SIX5. AP2A1 is identified as a gene that serves as a hub associated with the G-quadruplex binding FMR1 [8], while DMPK and SIX5 are suggested to be regulated by G4 motifs in two distinct studies [4, 9].

**Table 2.**
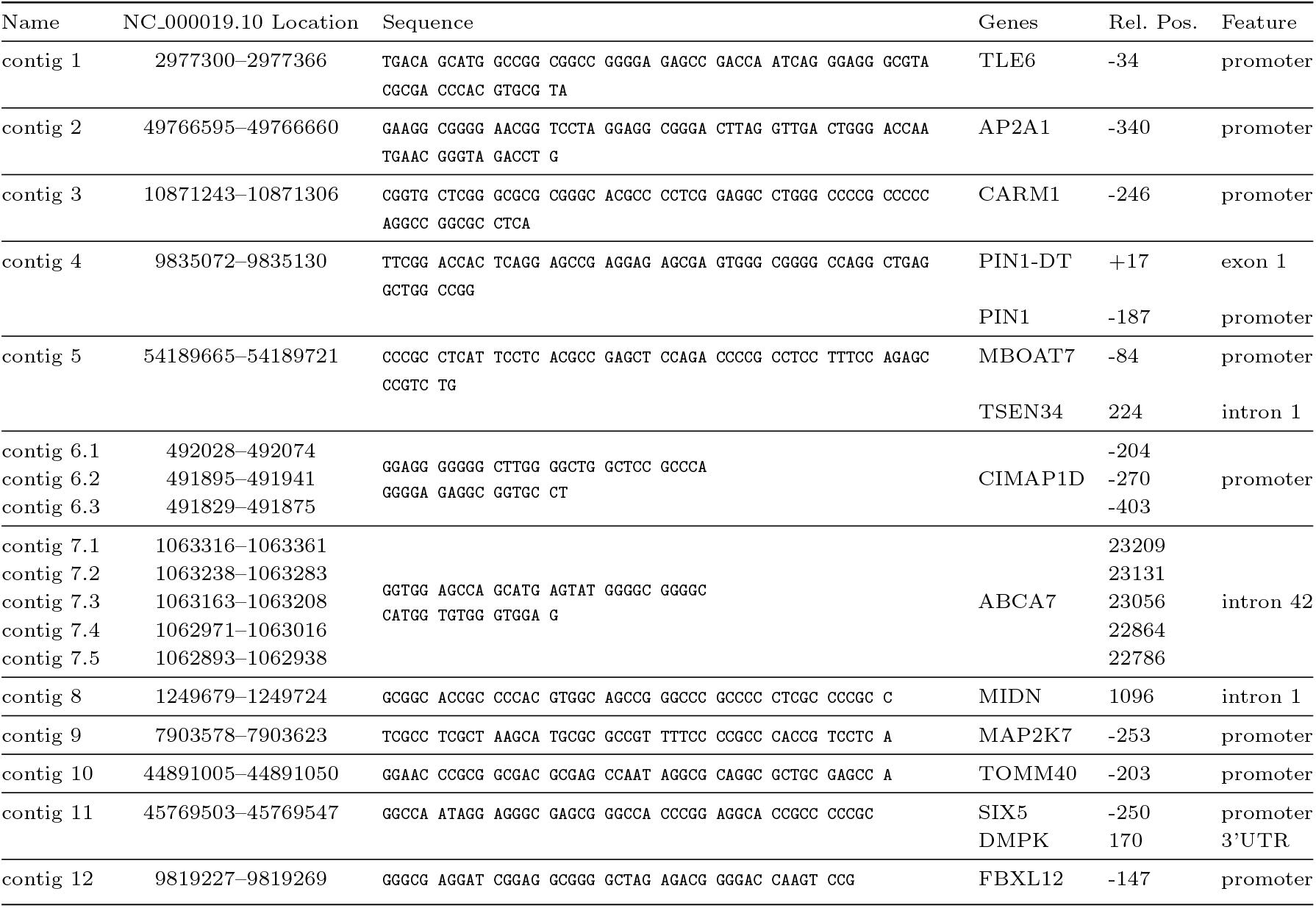
Contigs supported by 120 or more 21-mers as annotated on human chromosome 19, NC 000019.10.

## Conclusion

In this study, we introduced QuadStack, a deep learning framework specifically designed to predict *in vivo* BG4-binding G-quadruplex-associated sequences from genomic DNA. By incorporating two biologically motivated convolutional modules, G4Stack Convolution (G4SC) and Reverse Complement Convolution (RCC), the model captures both G/C-driven stacking patterns and strand-invariant features characteristic of ChIP-seq data. The combined architecture demonstrated cumulative improvement in performance, highlighting the value of embedding domain-specific inductive biases into deep learning models.

Using a large, multi-study BG4-ChIP-seq compendium, QuadStack achieved strong predictive performance and substantially outperformed existing *in vitro*-oriented predictors on independent genomic test data. These results underscore a fundamental discrepancy between *in vitro* G-quadruplex folding propensity and *in vivo* BG4-binding, suggesting that additional sequence and contextual constraints govern G4 behavior in the cellular environment.

Our results indicate that BG4-binding sequences extend beyond canonical G-rich motifs and involve mixed G/C patterns. This challenges the assumption that cytosines are uniformly disruptive and suggests a context-dependent role for cytosines *in vivo*. Alternatively, potential contributions from complementary-strand guanines to G-tetrad formation may also provide a plausible explanation for the differences between *in vivo* BG4 binding and canonical *in vitro* G4 rules. Future studies integrating structural and genomic approaches will be essential to resolve the underlying mechanisms.

Overall, QuadStack provides a robust computational framework for studying *in vivo* G-quadruplex-associated signals and offers a foundation for future investigations integrating chromatin context, epigenetic features, and structural constraints to further refine our understanding of G4 biology in genomic settings.

## Data availability

All code used for model development, training, and analysis is publicly available at https://github.com/odoluca/QuadStackML.

## Author contributions statement

P.N.U. curated the data, established and performed galaxy workflow, conceived the experiments. O.D. wrote the machine learning scripts and analyzed the results. P.N.U. and O.D. wrote and reviewed the manuscript.

## Acknowledgments

The authors thank the reviewers for their valuable suggestions.

## Osman Doluca

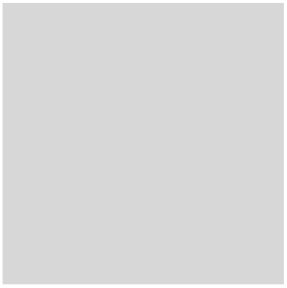

Dr. Doluca is an Associate Professor in the Department of Biomedical Engineering at the Izmir University of Economics, where he also serves as the lead investigator of the Doluca Lab. He earned his PhD in Biochemistry from Massey University, New Zealand, specializing in the structural dynamics of nucleic acids.

His research focuses on the intersection of bionanotechnology and computational biology, with a particular emphasis on the characterization of non-canonical DNA structures, such as G-quadruplexes, and their applications in biosensing and gene regulation.

## Polen Nehir Ulaş

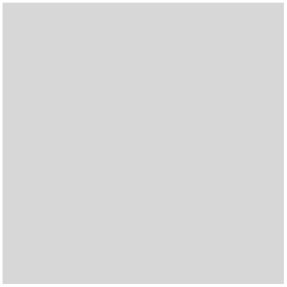

Ulaş received her BSc degree in Molecular Biology and Genetics from Izmir Institute of Technology and her MSc degree in Bioengineering from Izmir University of Economics. Her research focuses on computational genomics, bioinformatics, and the application of machine learning methods to genomic data.

